# Viral interference between severe acute respiratory syndrome coronavirus 2 and influenza A viruses

**DOI:** 10.1101/2024.02.02.578538

**Authors:** Shella Gilbert-Girard, Jocelyne Piret, Julie Carbonneau, Mathilde Hénaut, Nathalie Goyette, Guy Boivin

## Abstract

Some respiratory viruses can cause a viral interference through the activation of the interferon (IFN) pathway that reduces the replication of another virus. Epidemiological studies of coinfections between SARS-CoV-2 and other respiratory viruses have been hampered by non-pharmaceutical measures applied to mitigate the spread of SARS-CoV-2 during the COVID-19 pandemic. With the ease of these interventions, SARS-CoV-2 and influenza A viruses can now co-circulate. It is thus of prime importance to characterize their interactions. In this work, we investigated viral interference effects between an Omicron variant and a contemporary influenza A/H3N2 strain, in comparison with an ancestral SARS-CoV-2 strain and the 2009 pandemic influenza A/H1N1 virus. We infected nasal human airway epitheliums with SARS-CoV-2 and influenza, either simultaneously or 24 h apart. Viral load was measured by RT-qPCR and IFN-α/β/λ1/λ2 proteins were quantified by immunoassay. Expression of four interferon-stimulated genes (ISGs; OAS1/IFITM3/ISG15/MxA) was also measured by RT-droplet digital PCR. Additionally, susceptibility of each virus to IFN-α/β/λ2 recombinant proteins was determined. Our results showed that influenza A, and especially A/H3N2, interfered with both SARS-CoV-2 viruses, but that SARS-CoV-2 only interfered with A/H1N1. Consistently with these results, influenza, and particularly the A/H3N2 strain, caused a higher production of IFN proteins and expression of ISGs than SARS-CoV-2. The IFN production induced by SARS-CoV-2 was marginal and its presence during coinfections with influenza was associated with a reduced IFN response. All viruses were susceptible to exogenous IFNs, with the ancestral SARS-CoV-2 and Omicron being less susceptible to type I and type III IFNs, respectively. Thus, influenza A causes a viral interference towards SARS-CoV-2 most likely through an IFN response. The opposite is not necessarily true, and a concurrent infection with both viruses leads to a lower IFN response. Taken together, these results help us to understand how SARS-CoV-2 interacts with another major respiratory pathogen.

**Author summary:** During the COVID-19 pandemic, non-pharmaceutical measures were able to reduce the spread of SARS-CoV-2 and most respiratory viruses. Since the ease of these measures, SARS-CoV-2 variants and other viruses, such as influenza A, have started to co-circulate and can now infect a same host and interact with each other. These interactions can lead to attenuated or aggravated infections and can affect the timing of epidemics. Therefore, it is very important to elucidate how the new SARS-CoV-2 interacts with other viruses to better predict their implications in human health and their epidemic activity. Our work contributes to better understand these interactions using viruses that have likely co-circulated after lifting mitigation interventions, i.e., SARS-CoV-2 Omicron variant and a contemporary influenza A/H3N2 strain. We studied how each virus may affect the other virus’ growth and how these interactions were associated with the innate immune response of the host. We found that a prior infection with influenza A can decrease the growth of SARS-CoV-2 while the latter reduces the innate immune response. Our results help to understand the interplay between SARS-CoV-2 and influenza A in the host and may improve mathematical models predicting epidemics.

## Introduction

Different respiratory viruses can infect the same host concurrently or sequentially and may thus interact with each other. The interaction can be either positive (additive or synergistic), negative (antagonistic) or neutral. Positive/negative interactions may result in an increased/decreased host susceptibility to infection by the second virus, viral loads and duration of viral shedding. In turn, these parameters may influence the rate of viral transmission at the population level. Viral interference represents a negative interaction where an infection by a first virus inhibits the infection of a second virus through the induction of a non-specific innate immune response [1]. Upon recognition of viral components, host cells produce type I interferons (IFNs; IFN-α/β) as well as type III IFNs (IFN-λ); the latter being mainly found in epithelial cells of the gastrointestinal and respiratory tracts [2]. IFN proteins stimulate the production of additional IFN molecules and the expression of a multitude of interferon-stimulated genes (ISGs) in infected and neighbouring cells, amplifying the immune response. Many ISGs act as inhibitors of the viral replication, and ISG induction contributes to the establishment of an antiviral state within cells [3]. This may result in a refractory period during which infection of these cells by another homologous or heterologous virus is reduced. Viral interference effects have been observed between different respiratory viruses in *in vitro* and *in vivo* models [4–8]. The implication of the IFN response has been confirmed in most of these reports. Epidemiologic studies also suggested that negative interactions between viruses can affect epidemic curves at the population level [4, 9]. For instance, in 2009, a human rhinovirus (HRV) epidemic peak was associated with a delay in the spread of the pandemic A/H1N1 influenza virus in different countries [10, 11].

During the coronavirus disease 2019 (COVID-19) pandemic, caused by the severe acute respiratory syndrome coronavirus 2 (SARS-CoV-2), the detection of many seasonal respiratory viruses dramatically decreased [12, 13], with the exception of some non-enveloped viruses such as rhinoviruses and adenoviruses. This was mainly due to the implementation in most countries of non-pharmaceutical interventions, including social distancing, the use of facemasks, hand sanitizing, isolation, and quarantine, to reduce the transmission of SARS-CoV-2 [14–18]. This caused a disturbance in seasonal epidemics and resulted in off-season resurgence of some viruses when SARS-CoV-2 mitigation measures were subsequently lifted [12, 19].

Some studies investigated the risk of coinfections with SARS-CoV-2 and other respiratory viruses at the onset of the pandemic, i.e., before implementation of non-specific measures. Stowe *et al.* observed that the risk of being infected with SARS-CoV-2 was 58% lower in influenza A-positive patients [20]. Furthermore, Nenna *et al.* reported that an early 2021 autumnal respiratory syncytial virus epidemic seemed to have been interrupted by the arrival of the new Omicron variant in the population [21]. However, conclusions about viral interference effects between SARS-CoV-2 and other respiratory viruses were difficult to establish at that time due to limited coinfection events in the population during the pandemic. This underlines the necessity for additional research work in order to better understand the interactions between SARS-CoV-2 and other respiratory viruses, especially with the influenza A virus (IAV), which may have a major impact on morbidity and mortality.

So far, studies evaluating viral interference effects between SARS-CoV-2 and other respiratory viruses have focused on the ancestral D614G mutant and early variants (e.g., Alpha, Beta, Delta). For instance, many investigators showed that influenza A and HRV interferes with SARS-CoV-2 [22–28]. However, most of these studies investigated potential viral interference events between viruses that did not have much opportunity to interact with SARS-CoV-2 during the pandemic, such as influenza A/H1N1pdm09-derived virus, which had nearly disappeared during SARS-CoV-2’s first pandemic wave in 2020 [29]. After easing the non-specific interventions in spring of 2022, a late epidemic of A/H3N2 virus was observed in North America while A/H1N1 circulation remained low [29]. At that time, the SARS-CoV-2 Omicron variant was highly prevalent [30], and interactions between these two viruses are most likely to have occurred.

In this paper, we investigated potential viral interference effects between clinical isolates of a 2022 influenza A/H3N2 strain and a contemporary SARS-CoV-2 variant, i.e., Omicron (B.A.1) using nasal reconstituted human airway epitheliums (HAEs) cultured at the air-liquid interface. As our group already showed the occurrence of viral interference between the ancestral SARS-CoV-2 D614G strain and the influenza A/H1N1pdm09 virus in the same experimental model [22], we compared both pairs of viruses to evaluate potential changes that may have arisen since the onset of pandemic. We found that a first infection with A/H3N2 strongly interfered with both Omicron and the ancestral D614G SARS-CoV-2 virus, while the opposite was not true. On the other hand, we observed that Omicron, and to a lesser extent the ancestral virus, interfered with A/H1N1. A/H1N1 also interfered with both SARS-CoV-2 viruses, but not as markedly as A/H3N2. We then evaluated the primary and secondary IFN responses during coinfections, as well as the susceptibility of each virus to exogenous type I and type III IFNs. Our results suggest that influenza A/H3N2 interferes with SARS-CoV-2 through an important IFN response. One the other hand, SARS-CoV-2 inhibits the IFN response during coinfections with IAV, which may reduce its ability to cause viral interference.

## Materials and methods

### Cells and viruses

VeroE6 cells (green monkey kidney) were purchased from the American Type Culture Collection (CRL-1586; Manassas, VA, USA). VeroE6/TMPRSS2 cells were provided by the NIBSC Research Reagent Repository (UK), with thanks to Dr. Makoto Takeda (University of Tokyo). ST6-Gal-I MDCK (Madin-Darby Canine Kidney) cells overexpressing the ⍺2-6 sialic acid receptor (MDCK ⍺2-6) were obtained from Dr. Y. Kawaoka (University of Wisconsin, Madison, WI, USA) [31]. VeroE6 and MDCK ⍺2-6 cells were cultured in minimum essential medium (MEM; Invitrogen, Carlsbad, CA, USA) supplemented with 10% fetal bovine serum (FBS; Invitrogen) and 1% HEPES. Culture medium for MDCK ⍺2-6 also contained puromycin (7.5 µg/ml). VeroE6/TMPRSS2 were cultured in Dulbecco’s modified Eagle’s medium (DMEM) supplemented with 10% FBS, 1% HEPES and 1 mg/ml geneticin (Life Technologies, Carlsbad, CA, USA). Nasal reconstituted HAEs (MucilAir™, pool of donors, EP02MP) and their culture medium were provided by Epithelix Sàrl (Geneva, Switzerland). HAEs were cultured in 24-well inserts at the air-liquid interface. All cells and HAE inserts were maintained at 37°C with 5% CO_2_.

Influenza A/H3N2 virus (clade 3C.2a1b.2a.2; isolated from a clinical sample collected in April 2022 in Quebec City, Canada) and influenza A/California/7/2009 H1N1pdm09 virus were amplified on MDCK α2-6 in MEM supplemented with 1% HEPES and 1 μg/ml trypsin treated with N-tosyl-L-phenylalanine chloromethyl ketone (Sigma, Oakville, ON, Canada). Viral titers were determined by plaque assays. SARS-CoV-2 strain Quebec/CHUL/21697, an ancestral strain bearing the spike substitution D614G (referred to as D614G), and SARS-CoV-2 strain Quebec/CHUL/904,274 (Omicron: B.1.1.529, sub-lineage BA.1.15; referred to as Omicron) were isolated from nasopharyngeal swabs recovered in Quebec City, Canada, in March 2020 and December 2021, respectively. D614G was amplified on VeroE6 cells in MEM supplemented with 1% HEPES. Omicron was amplified on VeroE6/TMPRSS2 cells in DMEM with 1% HEPES. Viral titers were then determined by plaque assays. All experimental work using infectious SARS-CoV-2 was performed in a Biosafety Level 3 (BSL3) facility at the CHU de Québec-Université Laval.

### Infection kinetics in HAEs

Before infection, the apical poles of HAEs were washed with 200 µl of pre-warmed Opti-MEM (Gibco; ThermoFisher Scientific, Waltham, MA, USA) during 10 min at 37°C and pipetting up and down a few times to remove the mucus layer. The apical poles of HAEs were infected with each single virus at a multiplicity of infection (MOI) of 0.02 (considering that each HAE was made of 500 000 cells), in 200 µl of Opti-MEM. HAEs were incubated for 1 h at 37°C with 5% CO_2_, and the inoculum was then removed. For simultaneous coinfections, both viruses, at the same MOI, were added in 200 µl of medium. Sequential coinfections were made with each viral infection occurring 24 h apart. The trans-epithelial electrical resistance (TEER) was measured every 48 h from the first infection day, using a Millicell^®^ ERS-2 Voltohmmeter (Sigma-Aldrich, St. Louis, MO, USA). In single infections and when specified, the viability of HAEs was assessed by a MTS assay (CellTiter 96^®^ AQueous One Solution Cell Proliferation Assay; Promega, Madison, WI, USA) by addition of 20 µl of MTS solution with 180 µl of Opti-MEM on the apical pole of HAEs. After an incubation of 1 h in the dark at 37°C with 5% CO_2_, the absorbance was measured at 490 nm in a 96-well plate, using a Synergy HTX multi-mode reader (BioTek Instruments, Winooski, VT, USA). Fig 1 summarizes the experimental design.

**Fig 1.**
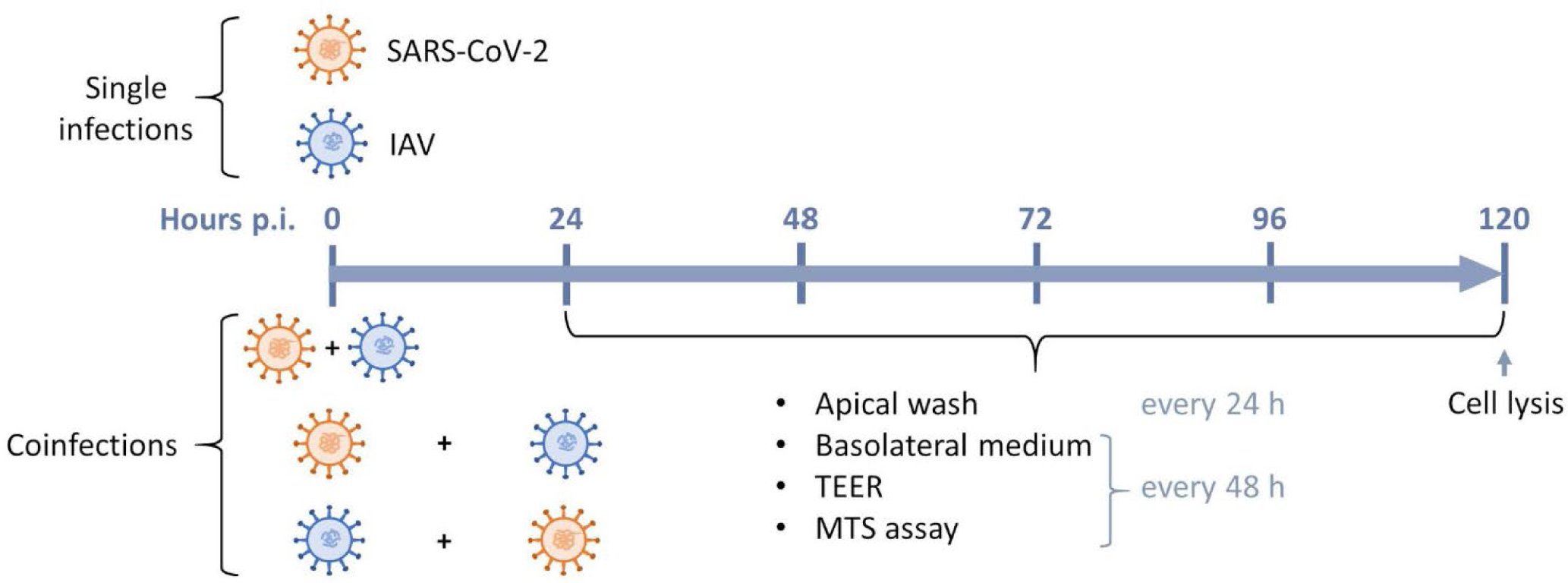
Timeline of SARS-CoV-2 and influenza A (IAV) single infections and coinfections in nasal human airway epitheliums (HAEs). In coinfection experiments, SARS-CoV-2 (orange) and influenza A (blue) were added simultaneously or 24 h apart at the apical pole of HAEs. Infections were monitored for 120 h after adding the first virus. Apical washes were collected every 24 h, while basolateral medium was taken and replaced by 500 µl of fresh media every 48 h. Trans-epithelial electrical resistance (TEER) was measured every 48 h starting from the day of first infection. MTS assays were performed every 48 h starting from the day before infection. HAEs were lysed with RNA extraction buffer at 120 h post-infection (p.i.).

Each day following the first infection, the apical pole of HAEs was washed with 200 µl of pre-warmed Opti-MEM by incubating for 10 min at 37°C with 5% CO_2_ and then pipetting the supernatant up and down. The apical wash was collected and stored at −80°C for viral RNA load determination. Every 48 h, the basolateral medium was taken and replaced with 500 µl of fresh pre-warmed MucilAir^TM^ culture medium. The collected basolateral medium was snap-frozen and stored at −80°C for cytokine quantification. After 120 h post-infection (p.i.), HAEs were lysed for ISG quantification by RT-droplet digital PCR. Uninfected HAEs were manipulated the same way as infected ones for 120 h prior to lysis.

### Treatment of HAEs with recombinant IFN proteins and IFN inhibitors

Recombinant human IFN-β (8499-IF) and IFN-λ2 (1587-IL) were purchased from R&D Systems (Minneapolis, MN, USA). Both were reconstituted in phosphate-buffered saline (PBS) with 0.1% bovine serum albumin (BSA; Sigma-Aldrich) and added to the basolateral pole at a concentration of 100 ng/ml. Recombinant human IFN-α2a (H6041; Sigma-Aldrich) was reconstituted in PBS with 0.1% BSA and added to the basolateral pole at a final concentration of 100 U/ml. HAEs were treated 24 h before primary infection and then daily in the basolateral medium until 120 h p.i.

Ruxolitinib (Cayman Chemical, Ann Arbor, MI, USA) was reconstituted in dimethyl sulfoxide (DMSO). It was added to the basolateral pole at a concentration of 5 µM 24 h prior to infection and was maintained at that concentration until 120 h p.i. BX795 (Sigma-Aldrich) was reconstituted in DMSO. It was added to the basolateral pole at a concentration of 6 µM 24 h prior to infection and was maintained at that concentration until 120 h p.i., as previously described in HAEs [4, 22, 23, 27].

### Viral RNA load quantification by RT-qPCR

Apical washes (100 µl) were first incubated in lysis buffer for 1 h at room temperature to inactivate SARS-CoV-2 before leaving the BSL3 facility. Viral RNA isolation was performed using the MagNA Pure LC system (Total nucleic acid isolation kit, Roche Molecular System, Laval, QC, Canada) or the EZ2 Connect system (EZ1&2 Virus Mini Kit v2.0, Qiagen, Toronto, ON, Canada). Then, reverse transcription quantitative PCR (RT-qPCR) assays were performed with the QuantiTect Virus + ROX Vial Kit (Qiagen, Toronto, ON, Canada) in a LightCycler^®^ 480 system (Roche Molecular System), using primers and probes targeting the M gene of influenza A (sequences available upon request) and the E gene of SARS-CoV-2 [32]. A value corresponding to the detection limit of the assays was attributed to samples with undetectable RNA levels.

### IFN protein quantification by magnetic bead-based immunoassay

Medium samples collected at the basolateral pole of HAEs (250 µl) were thawed and inactivated with 1% Triton X-100 for 1 h at room temperature before leaving the BSL3 facility. A multiplex magnetic bead-based immunoassay was performed for four targets (IFN-α, IFN-β, IL-28A/IFN-λ2 and IL-29/IFN-λ1) using a Bio-Plex Pro^TM^ Human Inflammation Panel 1 Express assay (Bio-Rad Laboratories Ltd., Mississauga, ON, Canada) according to the manufacturer’s instructions. Mean fluorescence intensity from all the bead combinations was measured using a Bioplex 200 system and the Bioplex Manager Software V6.2 (Bio-Rad Laboratories Ltd.).

### ISG expression by RT-ddPCR

After 120 h p.i., HAEs were treated with 100 µl of lysis buffer for 1 h at room temperature to inactivate SARS-CoV-2 before leaving the BSL3 facility. Viral RNA isolation was performed using the MagNA Pure LC system (Total nucleic acid isolation kit, Roche Molecular System) or the EZ2 Connect system (EZ1&2 Virus Mini Kit v2.0, Qiagen). Then, one-step reverse transcription droplet digital PCR (RT-ddPCR) assays were performed with the One-step RT-ddPCR Advanced Kit for probes (Bio-Rad Laboratories Ltd.), using primers and probes targeting 2’,5’-oligoadenylate synthetase 1 (OAS1), interferon-induced transmembrane protein 3 (IFITM3), interferon-stimulated gene 15 (ISG15), and myxovirus resistance protein A (MxA). Primers and probes are described in S1 Table. Expression of the ISGs was compared to that of a housekeeping gene (18S). For this latter gene, reverse transcription was done separately using the SuperScript™ IV First-Strand Synthesis System (Invitrogen™, ThermoFisher Scientific) according to the manufacturer’s instructions using 5 µl of RNA. The ddPCR reaction was performed using QX200™ ddPCR™ EvaGreen SuperMix (Bio-Rad Laboratories Ltd.). For all ddPCR experiments, droplets were generated using a QX200™ Droplet Generator (Bio-Rad Laboratories Ltd.) and PCR reactions were performed using a C1000 Touch Thermal cycler (Bio-Rad). Acquisition was made with a QX200™ Droplet Reader (Bio-Rad Laboratories Ltd.), with the software QX Manager 1.2.

### Statistical analysis

Statistical analyses were performed with GraphPad Prism version 9.4.0 (GraphPad Software, La Jolla, CA, USA). A one-way Brown-Forsythe and Welch analysis of variance (ANOVA) test with posthoc Dunnett’s T3 multiple comparisons test was used to compare viral RNA loads, ISG mRNAs or IFN protein levels in the different experimental conditions.

## Results

### Interference between SARS-CoV-2 and IAV

We first investigated the interactions between the contemporary SARS-CoV-2 Omicron variant and the influenza A/H3N2 strain that were circulating after the ease of non-pharmacological interventions. Reconstituted HAEs were infected with each single virus or coinfected with the two viruses simultaneously or sequentially (24 h apart). Fig 2A shows that a prior infection of HAEs with A/H3N2 greatly reduced the replication of Omicron by 3 logs compared to Omicron alone at 96 h p.i. However, when HAEs were infected with Omicron first or simultaneously to A/H3N2, the replication of Omicron was similar to that of the single infection. Fig 2B shows that in all A/H3N2 and Omicron coinfections (either simultaneous or sequential), the growth of A/H3N2 was comparable to that of the single virus. Thus, although A/H3N2 interferes with Omicron when added 24 h earlier, the opposite is not true. When investigating the interactions between the SARS-CoV-2 and the IAV subtype circulating at the onset of the pandemic (i.e., in winter 2020), our group previously showed that ancestral D614G interfered with influenza A/H1N1pdm09 virus [22]. We thus evaluated whether, in contrast to Omicron, SARS-CoV-2 D614G would interfere with an A/H3N2 strain from 2022 in HAEs (Fig 2C-D). With this strain as well, we observed that A/H3N2 causes viral interference towards SARS-CoV-2, but not the opposite.

**Fig 2.**
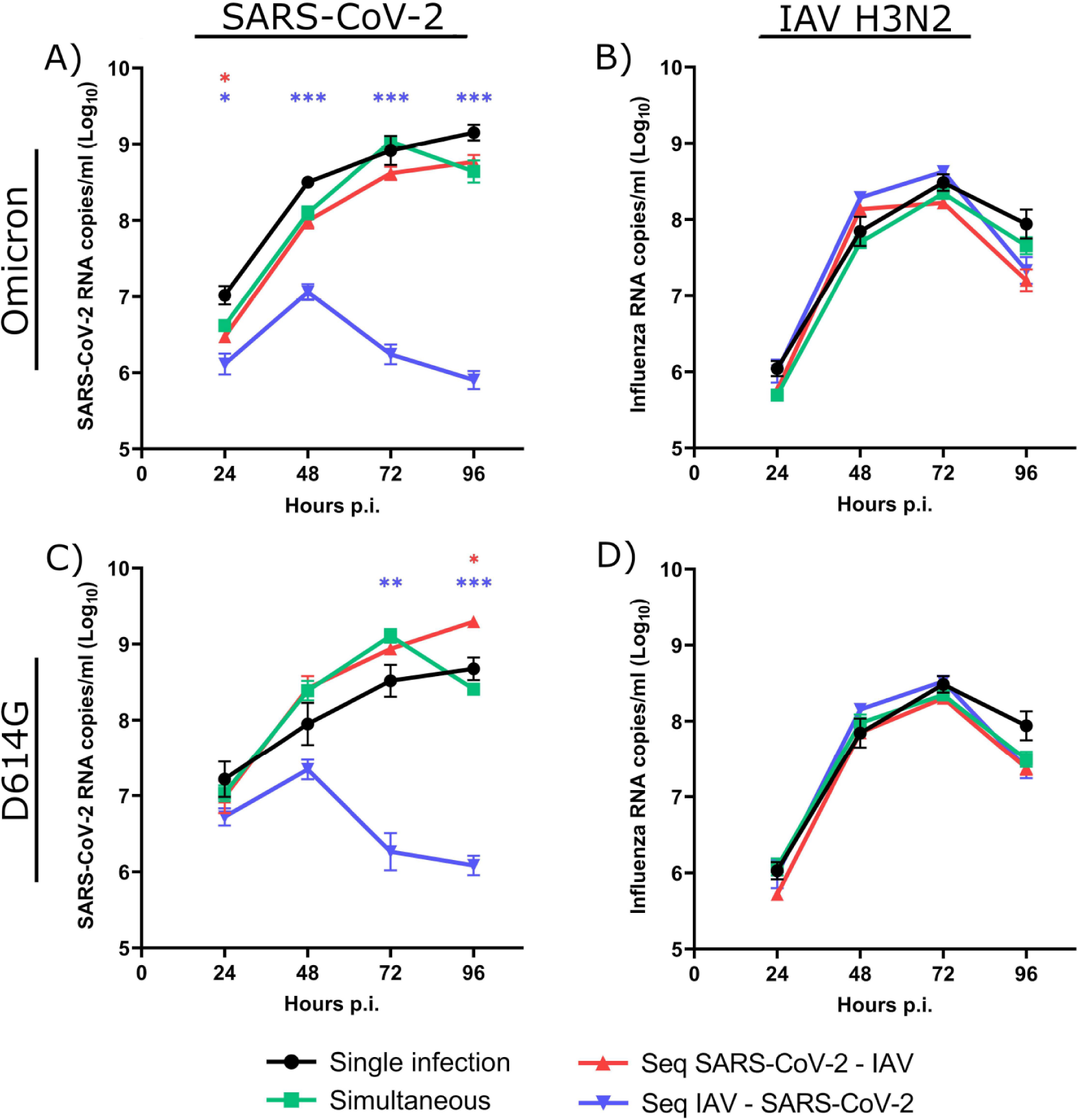
Viral interference between SARS-CoV-2 strains and influenza A/H3N2. Viral RNA loads of SARS-CoV-2 (Omicron variant (A) or D614G mutant (C)) and influenza A/H3N2 (B, D) during single infections or simultaneous and sequential (seq) coinfections in nasal human airway epitheliums (HAEs). Hours post-infection (p.i.) represent the time after the infection with either SARS-CoV-2 (A, C) or influenza (B, D). Results are expressed as the mean of the Log_10_ of viral RNA copies per ml ± SEM of 3-4 replicates of HAEs in one experiment. *: p ≤ 0.05, **: p ≤ 0.01, ***: p ≤ 0.001. Color of asterisks corresponds to that of the curve, compared to the single infection.

We next tested coinfections between the two SARS-CoV-2 strains and A/H1N1. When A/H1N1 was the primary virus, the growth of Omicron was reduced by over 1 log throughout the infection (Fig 3A). However, in contrast to A/H3N2, in HAEs infected with Omicron first, the growth of A/H1N1 was inhibited (Fig 3B). Interestingly, in HAEs infected with A/H1N1 prior to Omicron, A/H1N1 seemed to grow faster than in other conditions. In sequential A/H1N1 and D614G coinfections (Fig 3C-D), A/H1N1 caused more interference than with Omicron since the viral load of D614G barely increased and was reduced by 2 logs at 96 h p.i., compared to D614G alone. The simultaneous coinfection of D614G and A/H1N1 also resulted in inhibition of SARS-CoV-2 throughout the infection, albeit not significantly. On the other hand, a 1-log reduction of A/H1N1 was observed from 48 h to 96 h p.i. when HAEs were infected with D614G first, although this was not statistically significant.

**Fig 3.**
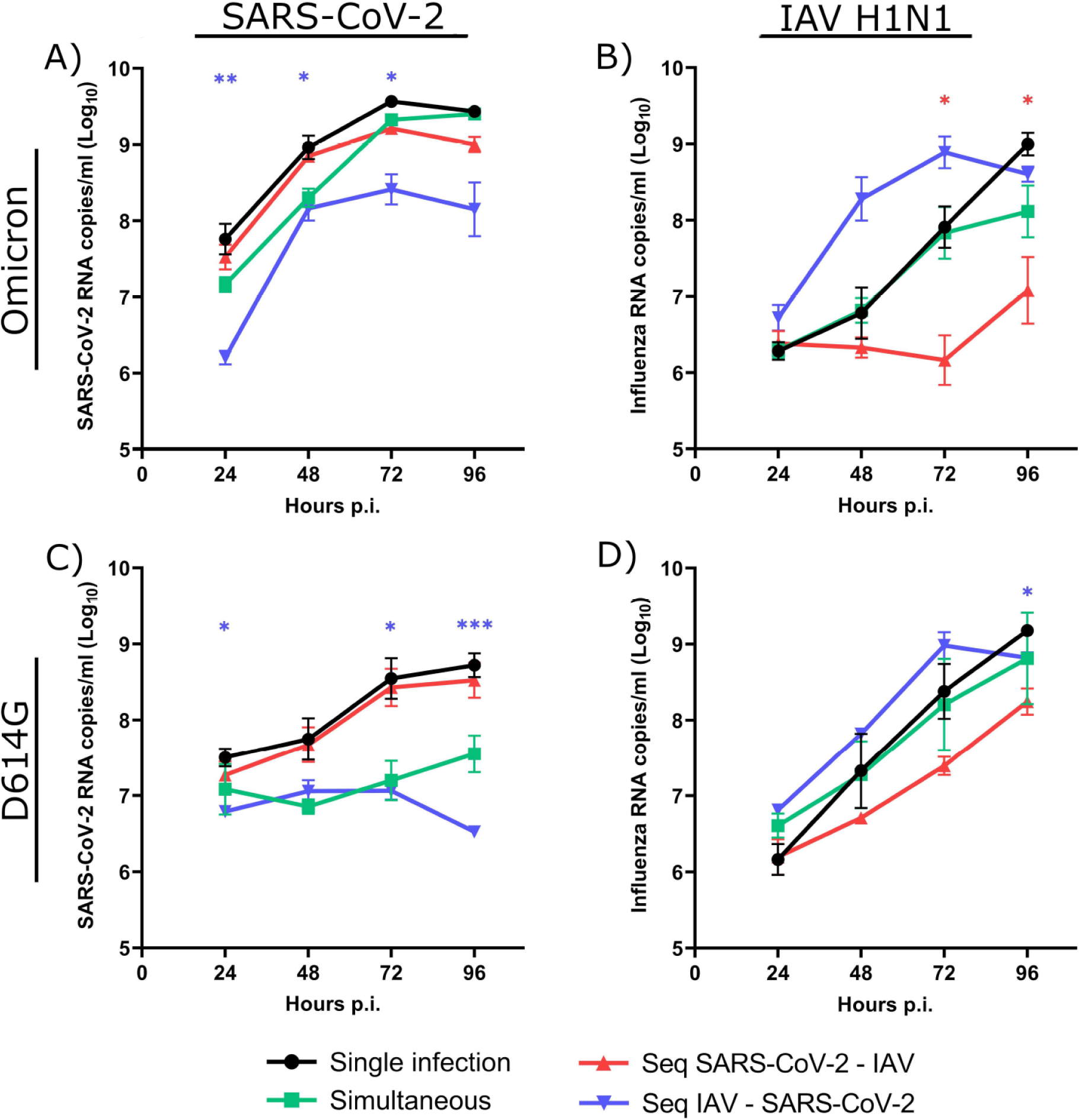
Viral interference between SARS-CoV-2 strains and influenza A/H1N1. Viral RNA loads of SARS-CoV-2 (Omicron variant (A) or D614G mutant (C)) and influenza A/H1N1 (B, D) during single infections or simultaneous and sequential (Seq) coinfections in nasal human airway epitheliums (HAEs). Hours post-infection (p.i.) represent the time after the infection with either SARS-CoV-2 (A, C) or influenza (B, D). Results are expressed as the mean of the Log_10_ of viral RNA copies per ml ± SEM of triplicate HAEs in one or two independent experiments. *: p ≤ 0.05, **: p ≤ 0.01, ***: p ≤ 0.001. Color of asterisks corresponds to that of the curve, compared to the single infection.

Of note, all viruses caused a reduction of the TEER of HAEs, especially A/H3N2 (panel A in S1 Fig). We thus verified the cellular viability of HAEs during single infections with each virus by MTS assays and concluded that all HAEs survived the infection (panel B in S1 Fig). Additionally, we measured the expression of a housekeeping gene (18S) at 120 h p.i. in lysates of infected HAEs and confirmed that cells still adhered to the insert membranes at the end of the kinetics experiment with all viruses (panel C in S1 Fig). Taken together, these results confirmed that HAEs infected with A/H3N2, albeit showing a higher reduction of the TEER than those infected with the other viruses, did survive until the end of the experiments. Thus, our results suggest that a primary infection with IAV interferes with SARS-CoV-2 D614G, and that the interference induced by A/H3N2 was more important than that of A/H1N1. However, SARS-CoV-2 Omicron, and to a lesser extent D614G, interfere only with A/H1N1.

### SARS-CoV-2 induces a weaker IFN response than IAV and inhibits IFN production in coinfections

To better understand the role of IFN in the viral interference process between IAV and SARS-CoV-2, we first investigated the production of type I and type III IFNs induced by viruses in single infections and coinfections. The basolateral medium of infected HAEs was collected every 48 h and the levels of IFN proteins (IFN-α/β/λ1/λ2) were measured by magnetic bead-based immunoassay. No IFN-α protein was detected in any condition, as reported elsewhere [25, 33]. At 24 h p.i., there was no IFN-β detected, and the production of IFN-λ1 and λ2 was minimal and not significantly different for all infection conditions tested (S2 Fig). IAV, especially A/H3N2, caused a much greater secretion of type I and type III IFN proteins than SARS-CoV-2, with maximal values reached at 72 h and 120 h p.i. for A/H3N2 and A/H1N1, respectively (Fig 4). Levels of type III IFNs, especially IFN-λ2, were much higher than those of IFN-β. Single infections with both SARS-CoV-2 viruses induced no IFN-β and only a marginal IFN-λ1 and λ2 production. Interestingly, IFN secretion was decreased in almost all coinfections with IAV compared to IAV alone, especially when SARS-CoV-2 was added first or simultaneously. This effect may result from the mechanisms of immune evasion induced by SARS-CoV-2 to escape or reduce IFN response [34, 35]. However, at 72 h p.i., levels of IFNs were more elevated in the sequential A/H1N1 – Omicron coinfection than with A/H1N1 in single infection or in other coinfection conditions (Fig 4G-H-I). We observed that A/H1N1 grew more quickly when it is added before Omicron (Fig 3), which may have led to a faster IFN response. Overall, the higher production of type I and type III IFNs induced by IAV, especially A/H3N2, could explain why it interferes more readily with SARS-CoV-2 compared to A/H1N1.

**Fig 4.**
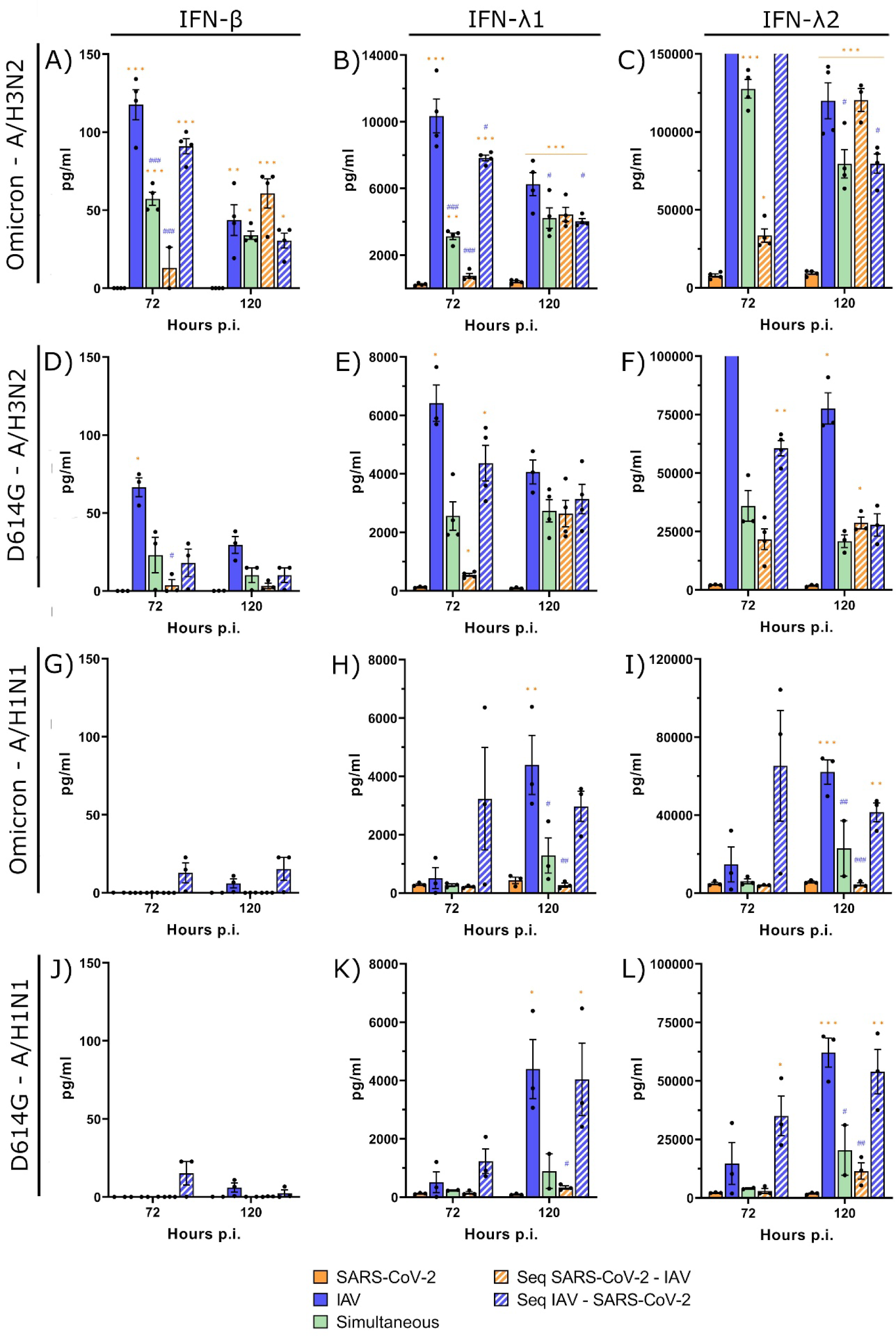
Interferon protein production during SARS-CoV-2 and influenza A coinfections. Type I and type III interferon (IFN-β, IFN-λ1, IFN-λ2) proteins production during single and coinfections of nasal human airway epitheliums with SARS-CoV-2 (Omicron and D614G) and influenza A (H3N2 and H1N1) at 72 h and 120 h post-infection (p.i.). Results are expressed as the mean amount of IFN proteins in pg per ml ± SEM of 2 to 6 replicates in one or two independent experiments (bars that appear higher than the Y axis maximum represent values that are “out of range”). Orange *: compared with Omicron alone, blue #: compared with A/H3N2 alone. *, #: p ≤ 0.05, **, ##: p ≤ 0.01, ***, ###: p ≤ 0.001.

### SARS-CoV-2 induces a weaker ISG expression than influenza A

Type I and type III IFNs are associated with the expression of several ISGs [2]. We thus investigated the expression of four ISGs acting on different steps of viral infection (i.e., OAS1, IFITM3, ISG15, MxA) in lysates of HAEs infected with A/H3N2, A/H1N1, D614G and Omicron viruses in single and coinfections. Uninfected HAEs exhibited a minimal ISG expression (panel A in S3 Fig) that was significantly lower than those of all infected HAEs (p ≤ 0.05). The expression of the different ISGs was almost similar between single infections with the two SARS-CoV-2 strains as well as between single infections with the two influenza A viruses (S3 Fig).

Fig 5 shows that SARS-CoV-2 (Omicron and D614G) induced a significantly lower expression (p ≤ 0.05) of the different ISGs than influenza A/H1N1 and A/H3N2. In all SARS-CoV-2 and A/H3N2 coinfections, the ISG expression was almost comparable to that of SARS-CoV-2 alone, regardless of the first infecting virus. In Omicron and A/H1N1 coinfections, all four ISGs were more inhibited when Omicron was the first virus. In the simultaneous coinfection or when A/H1N1 was the primary virus, expression of the ISGs was more often intermediate between that of Omicron and A/H1N1. In contrast, in D614G and A/H1N1 coinfections, the expression of almost all ISGs was more or less similar to that induced by A/H1N1 alone. These results thus partly reflect what was observed with the primary IFN response, with a stronger immune response being induced by IAV than by SARS-CoV-2.

**Fig 5.**
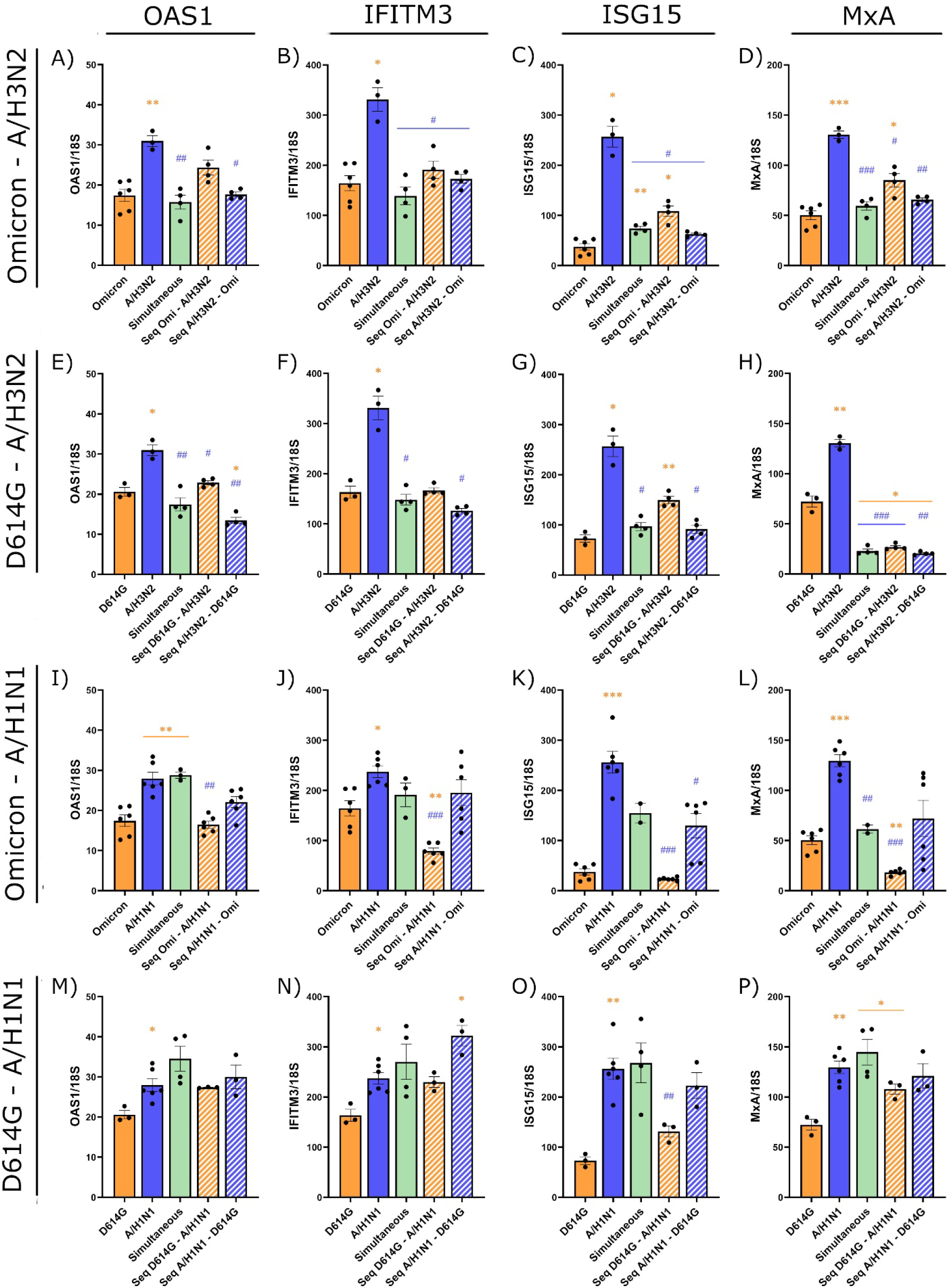
Interferon-stimulated gene expression during SARS-CoV-2 and influenza A coinfections. Expression of four interferon-stimulated gene (ISG) mRNAs (OAS1, IFITM3, ISG15, MxA) at 120 h post-infection during single infections or coinfections (simultaneous or sequential (Seq)) with IAV and SARS-CoV-2 strains. Nasal human airway epitheliums were infected with Omicron (Omi) and A/H3N2 (panels A-D), D614G and A/H3N2 (panels E-H), Omicron and A/H1N1 (panels I-L), and D614G and A/H1N1 (panels M-P). Results are expressed as the mean of the ratio of ISG mRNAs over that of 18S housekeeping gene (both in copies per µl) ± SEM of 3 to 6 replicates in one or two independent experiments. orange *: compared with Omicron alone, blue #: compared with A/H3N2 alone. *, #: p ≤ 0.05, **, ##: p ≤ 0.01, ***, ###: p ≤ 0.001.

### SARS-CoV-2 and IAV have similar susceptibility to type I and III IFNs

The susceptibility of different viruses to the IFN response is another factor that could affect viral interference effects [1]. We thus assessed the susceptibility of SARS-CoV-2 and IAV to exogenous type I and III IFNs by treating infected HAEs with recombinant IFN-α2a, −β and −λ2 proteins (Fig 6). The viral RNA load of SARS-CoV-2 Omicron was markedly decreased by 3 to 4 logs at 120 h p.i. in the presence of IFN-α and β. Treatment with type III IFN was much less effective, causing a 1-log reduction of the viral RNA load of Omicron early in the infection. The viral RNA load of Omicron eventually reached the same level as that of untreated controls later during the infection. D614G was slightly less susceptible than Omicron to IFN-α and β (difference not significant at most time points), with a reduction of its growth slightly lower than 3 logs, but it was a little more susceptible to IFN-λ2 (2-log reduction) at 120 h p.i. Both A/H3N2 and A/H1N1 were more susceptible to type I IFNs than SARS-CoV-2 (especially D614G), with about 4-log of reduction at 120 h p.i. There was no significant difference between the two IAVs but A/H3N2 was slightly less affected by IFN-λ2, with only 1-log reduction, while the growth of A/H1N1 was inhibited by 2 logs throughout the infection. Thus, the viral interference between SARS-CoV-2 and influenza does not appear to be related to a difference in their susceptibility to IFN.

**Fig 6.**
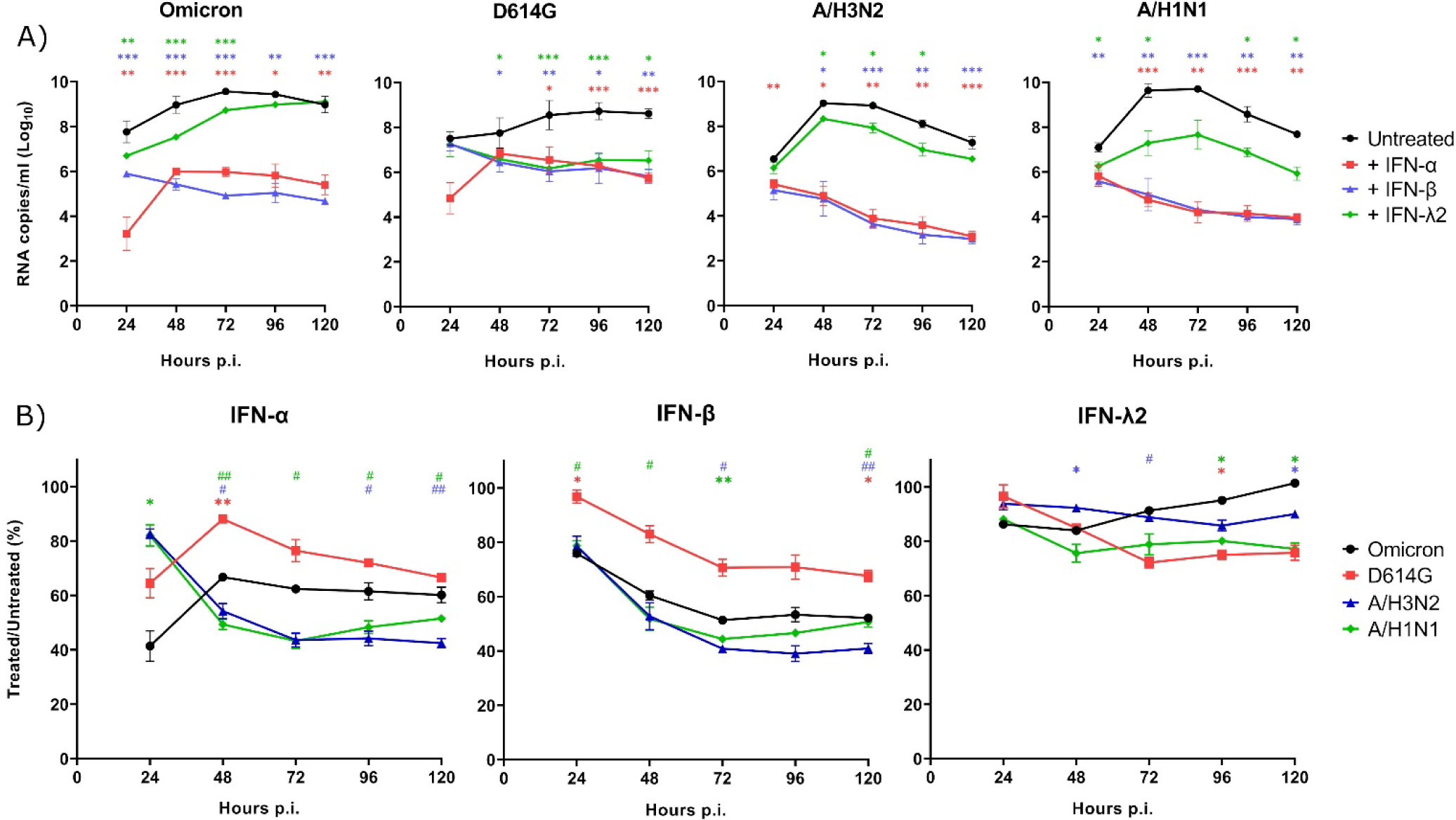
Susceptibility of SARS-CoV-2 and influenza A to recombinant interferon proteins. A) Viral RNA loads in nasal human airway epitheliums (HAEs) infected with SARS-CoV-2 (Omicron and D614G) and influenza (A/H3N2 and A/H1N1), in presence or absence of recombinant IFN-α2a, IFN-β or IFN-λ2. Results are expressed as the mean of the Log_10_ of viral RNA copies per ml ± SEM of triplicate HAEs in one experiment. A value of 60 copies/ml, corresponding to the detection limit of the assays, was attributed to samples with undetectable RNA levels (n=3). *: p ≤ 0.05, **: p ≤ 0.01, ***: p ≤ 0.001, in comparison with untreated HAEs. B) Comparison of the effects of IFN on the different viruses. Results are expressed as the mean percentage of the viral RNA loads of IFN-treated over untreated HAEs ± SEM using triplicates in one experiment. *: comparison with Omicron, #: comparison with D614G. *, #: p ≤ 0.05, **, ##: p ≤ 0.01. Color of symbols corresponds to that of the curve. No significant difference was observed between A/H3N2 and A/H1N1.

### Effects of an IFN inhibitor on viral replication and interference

Finally, we investigated whether viral interference would still occur in the presence of an IFN inhibitor. We used ruxolitinib, a JAK1-JAK2 inhibitor that has been approved for the treatment of multiple diseases, such as myelofibrosis, osteofibrosis, polycythemia vera, and steroid-refractory acute graft-*versus*-host disease. We first treated HAEs with ruxolitinib before and during single infections to evaluate its effects on the viral growth of IAV and SARS-CoV-2 (Fig 7). As expected, ruxolitinib increased the viral RNA loads of D614G and A/H1N1 by up to 1.5 and 2 logs, respectively. In contrast, the replication of Omicron and A/H3N2 remained mostly unaffected. We next looked at the effect of ruxolitinib on the viral interference between influenza and SARS-CoV-2 Omicron. However, we found that the growth of Omicron in coinfection with both IAV was still reduced in the presence of ruxolitinib (Fig 7C-D). Indeed, a prior A/H3N2 infection reduced by more than 3 logs the viral RNA load of Omicron in presence (Fig 7) or absence (Fig 2) of ruxolitinib, whereas a primary A/H1N1 infection decreased the viral RNA load of Omicron by 2.5 logs and 1.5 logs with (Fig 7) and without (Fig 3) ruxolitinib, respectively. Thus, the IFN inhibitor did not rescue the replication of Omicron during sequential coinfections with IAV.

**Fig 7.**
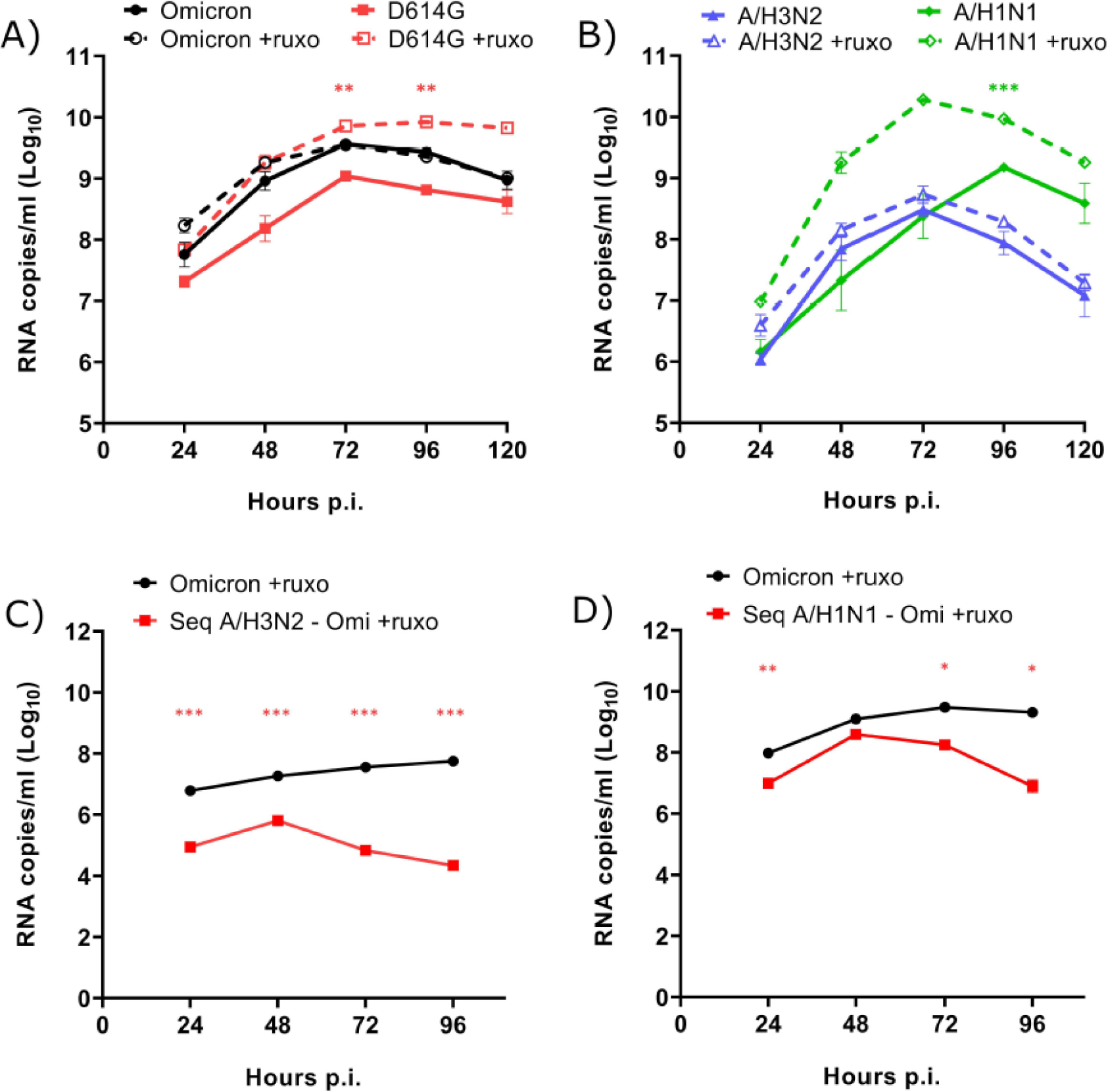
Effect of an interferon inhibitor on viral interference between SARS-CoV-2 Omicron and influenza A. Viral RNA loads of A) SARS-CoV-2 (Omicron and D614G) and B) influenza (A/H3N2 and A/H1N1) during single infections in nasal human airway epitheliums (HAEs), in presence or absence of ruxolitinib (ruxo). C-D) Effects of ruxolitinib on viral RNA loads of SARS-CoV-2 Omicron during single infections or sequential (seq) coinfections 24 h after (C) A/H3N2 and (D) A/H1N1. Results are expressed as the mean of the Log_10_ of viral RNA copies per ml ± SEM of 3-4 replicates in one experiment. *: p ≤ 0.05, **: p ≤ 0.01, ***: p ≤ 0.001. Color of asterisks corresponds to that of the curve, compared to untreated HAEs (A, B) or to the single infection (C, D).

However, we observed a marked drop of the TEER in all infected HAEs, especially when IAV infections were done in the presence of ruxolitinib (panel A in S4 Fig). The viability of HAEs infected with IAV and treated with ruxolitinib was also markedly decreased when assessed by MTS assays (especially for A/H3N2; panel B in S4 Fig) and 18S mRNA quantification at 120 h p.i. (for both A/H1N1 and A/H3N2; panel C in S4 Fig). Furthermore, an almost complete loss of cells on the inserts was seen at the end of the kinetics experiments. Nevertheless, we were able to confirm the inhibitory activity of ruxolitinib on ISG expression in HAEs infected with SARS-CoV-2, which were still viable at 120 h p.i. (panel D in S4 Fig).

We then evaluated the effects of another IFN inhibitor, BX795, which inhibits TANK-binding kinase 1 activity, on the viral interference between A/H3N2 and Omicron (S5 Fig). We also observed that the IFN inhibitor did not rescue the growth of Omicron and resulted in an even larger difference in viral RNA loads (almost 4 logs) compared to that of Omicron alone. With BX795 as well, infection with A/H3N2 resulted in a highly increased cell death rate, as indicated by TEER measurements and MTS assays (panels B-C in S5 Fig). Overall, our results suggest that a rescue of Omicron replication could not be seen due to the increased cell death in HAEs infected with IAV in the presence of both IFN inhibitors.

## Discussion

The COVID-19 pandemic has dramatically affected the circulation of seasonal viruses, mainly as a result of the introduction of non-pharmaceutical interventions to mitigate the spread of SARS-CoV-2. Nevertheless, viral interference events have been reported between SARS-CoV-2 and other respiratory viruses [20, 21]. Previous studies using human respiratory epitheliums have already shown viral interference effects between the ancestral SARS-CoV-2 (D614G) and influenza A/H1N1pdm09 (the subtype that circulated at the onset of the COVID-19 pandemic) [22, 24, 25, 36]. In this paper, we further explored events that have likely happened after lifting non-pharmacological measures by investigating the interactions between contemporary SARS-CoV-2 Omicron and IAV H3N2 (clade 3C.2a1b.2a.2) viruses. We also compared both pairs of viruses to evaluate whether potential changes in their interactions may have occurred over time. To better understand the role played by IFN in the viral interference effects, we looked at the primary and secondary IFN responses induced by each virus, as well as their susceptibility to type I and type III IFN proteins.

We found that both influenza A/H1N1 and A/H3N2 reduced the replication of SARS-CoV-2 D614G in a similar manner whereas A/H3N2 caused more interference with Omicron than A/H1N1. The two SARS-CoV-2 strains, and mainly Omicron, also interfered with A/H1N1, but not with A/H3N2. The main IFN proteins induced by IAV and SARS-CoV-2 infections were IFN-λ1 and λ2, which is in accordance with previous reports showing that type III IFNs are the first and predominant antiviral response in airway epitheliums [2, 37]. In agreement with previous reports [23, 25, 27, 33, 36], we observed that IAV strains caused a more important IFN-β and IFN-λ release in HAEs compared to SARS-CoV-2 strains. Among the two influenza strains, A/H3N2 induced the strongest IFN production. As the IFN response, the ISG expression was also higher during IAV infections than with SARS-CoV-2. All viruses were more susceptible to type I than to type III exogenous IFNs. Compared to Omicron, we observed that D614G exhibits a slightly lower susceptibility to IFN-I and a slightly higher susceptibility to IFN-λ2. These results partly differ from previous works showing that more recent SARS-CoV-2 variants were less susceptible than earlier strains to type III and type I IFNs [38, 39]. Differences in strains and cells used could account for this discrepancy.

During coinfections with A/H3N2, we observed that a first infection with the influenza virus strongly reduced the replication of both SARS-CoV-2 D614G and Omicron, while SARS-CoV-2 did not interfere with A/H3N2. Another study, using bronchial HAEs, showed that A/H3N2 interfered with SARS-CoV-2 Beta variant, but it was not affected by SARS-CoV-2 [23]. This is consistent with A/H3N2 inducing strong primary and secondary IFN responses, which can inhibit subsequent infection by another virus. In contrast, an infection with SARS-CoV-2, which does not lead to a strong activation of the IFN response, is less likely to cause viral interference. In this context, SARS-CoV-2 only induced the production of low amounts of type III IFN and A/H3N2 was not very susceptible to IFN-λ. Furthermore, it is well known that many proteins of SARS-CoV-2 can inhibit the primary and secondary IFN responses by targeting various components of the signaling pathways [34, 35, 40–44]. During our coinfection studies with A/H3N2, the mechanisms of immune evasion of SARS-CoV-2 seemed to have inhibited the IFN response, especially when SARS-CoV-2 was the primary virus or during simultaneous coinfections. Expression of all four ISGs (OAS1, IFITM3, ISG15, MxA) was also reduced in all coinfections with SARS-CoV-2 (either D614G or Omicron) and A/H3N2, compared to that of A/H3N2 alone. Regardless of which virus was infecting first, SARS-CoV-2 reduced ISG expression to levels similar to those induced by SARS-CoV-2 alone. Another factor that could be at play in the viral interference observed is that SARS-CoV-2 has a slower growth rate than IAV, which makes it more susceptible to interactions with faster growing viruses [45]. Furthermore, as A/H3N2 causes more damage to the HAEs than SARS-CoV-2, as reported by others [23], the number of host cells available for SARS-CoV-2 infection may be reduced, which may contribute to the interfering effect induced by A/H3N2.

We also confirmed results of previous reports showing that A/H1N1pdm09 interfered with ancestral SARS-CoV-2 in human respiratory epitheliums [22, 24, 25, 36] and extended these data to the Omicron variant. We observed that A/H1N1 did not interfere as strongly with Omicron as A/H3N2. This could be partly related to the weaker IFN response induced by A/H1N1 compared to A/H3N2. Based on our kinetics experiments, the growth rate of A/H1N1 was also slightly slower than that of A/H3N2 suggesting that A/H1N1 may have a lower ability to interfere with other viruses [45]. Furthermore, Omicron might have developed more effective mechanisms to evade the low IFN response caused by A/H1N1 than D614G, as suggested in some reports [38, 39, 46, 47]. Compared to D614G, Omicron was slightly more sensitive to IFN-α and −β but it was less sensitive to IFN-λ2, which was mainly expressed during IAV infection. Although IFN production was almost similar in coinfections with A/H1N1 and both SARS-CoV-2 viruses, ISG expression seemed to be higher during coinfections with D614G than with Omicron. This could also explain the stronger interference of A/H1N1 towards D614G compared to Omicron. Contrarily to what was observed with A/H3N2, SARS-CoV-2 Omicron interfered with A/H1N1. D614G showed a tendency to interfere with A/H1N1 as well, but this effect was not significant. We may suggest that although SARS-CoV-2 does not induce a strong IFN response, the low amount of IFN-λ produced might be sufficient to affect the growth of A/H1N1, which shows a tendency to be slightly more susceptible to type III IFN than A/H3N2. Nevertheless, the observation that SARS-CoV-2 could interfere with IAV is contradictory in several reports [22, 24–26, 36]. Differences in viral strains, host cells, timing of infections and study designs might explain these inconsistent results.

To confirm the involvement of the IFN response in viral interference effects, HAEs were incubated prior and during single and coinfections with an IFN inhibitor, ruxolitinib. During single infections with D614G and A/H1N1, viral RNA loads were increased in presence of ruxolitinib, as previously observed by our group with another IFN inhibitor, BX795 [22]. In contrast, ruxolitinib did not affect the growth of Omicron and A/H3N2. Shalamova *et al.* also observed that the growth of the ancestral SARS-CoV-2, but not Omicron, was increased in presence of ruxolitinib [46]. The effects of ruxolitinib and BX795 on the growth of IAV and SARS-CoV-2 reported in several papers [22, 23, 46, 48–50] were highly divergent; some described no significant effect whereas others showed an increased IAV and SARS-CoV-2 replication. The nature of these discrepancies could be related to the experimental conditions and viral strains used. Surprisingly, in our experiments, ruxolitinib and BX795 did not rescue the replication of Omicron in HAEs coinfected with A/H3N2 or A/H1N1. This lack of effects was related to the rapid and severe damage in HAEs infected with IAV in the presence of ruxolitinib or BX795. As we observed no severe damage in HAEs infected with SARS-CoV-2, we could confirm that ruxolitinib effectively inhibited the ISG expression. Although we did not evaluate the cytotoxic concentrations of both inhibitors in HAEs, concentrations of at least 5 µM of ruxolitinib and 6 µM of BX795 were not shown to cause any cytotoxicity in various cell lines, including human airway cells [51–56]. For instance, nasal HAEs have been exposed to 10 µM of ruxolitinib [50] and 6 µM of BX795 [4, 22, 23] without any cytotoxicity being noted. One possible explanation could be that combined effects between IAV infection and IFN inhibition may cause increased cell death in HAEs. Thus, IFN response inhibition with ruxolitinib or BX795 did not allow the rescue of Omicron during coinfections with IAV due to unexpected cell death, suggesting that other ways to block the IFN response should be envisaged (for instance, the use of antibodies).

In this paper, we compared viral interference effects between IAV and SARS-CoV-2 viruses that likely interacted at the onset of the pandemic and when lifting the non-pharmaceutical interventions that prevented their co-circulation. We used reconstituted human nasal epitheliums obtained from a pool of donors, which is a respiratory infection model more representative of clinical infections than cultured cell lines. This model allows to control the timing of infection (simultaneous or sequential) to study how viruses interact with each other as well as the primary and secondary IFN responses. In our study, the TEER measurement, which is generally used in experiments conducted in HAEs, did not seem to be a precise indicator of cell survival. We thus used complementary tests, such as the determination of the expression of a housekeeping gene (18S) and MTS assays. We suggest that combining these two assays with TEER measurements could be more appropriate to evaluate cell viability when using HAEs. Nonetheless, this study has some limitations as HAEs remain an incomplete model that does not account for the role of immune cells and other components of the adaptive immune system. Finally, the unexpected cell death observed in HAEs infected with IAV in the presence of IFN inhibitors may have prevented the rescue of Omicron replication. More research using these inhibitors or other ways to block the IFN response, such as antibodies, in presence of respiratory viruses will be needed to fully understand these observations.

## Conclusion

In this paper, we showed that IAV, and especially A/H3N2, interferes with SARS-CoV-2 Omicron, while Omicron interferes with A/H1N1 only. These results are in agreement with a recent retrospective study that showed a negative correlation between SARS-CoV-2 and influenza activity and an alternating dominance between the two viruses since the arrival of the Omicron variant [57]. All four viruses were shown to be sensitive to exogenous IFNs, especially to type I IFN response. Thus, the interfering effect of IAV on SARS-CoV-2 is probably due to the more potent primary and secondary IFN responses induced by IAV. SARS-CoV-2 demonstrated a tendency to inhibit IFN production and only induced a very limited IFN response. We cannot exclude, however, that other intrinsic virus-specific inhibition mechanisms could also be involved in these viral interference effects [58]. A better understanding of viral interference between respiratory viruses could help to improve mathematical models of viral transmission to predict epidemics and future pandemics and to make public health recommendations. New non-specific therapeutic avenues based on activation of the innate immune response for treatment of viral infections may also arise from this knowledge.

## Acknowledgments

We acknowledge the bioimaging platform of the Infectious Disease Research Centre, funded by an equipment and infrastructure grant from the Canadian Foundation Innovation.

## Supporting information captions

**S1 Table. Sequences of primers and probes used for quantification of interferon-stimulated genes and a housekeeping gene by ddPCR.**

**S1 Fig. Viability of nasal human airway epitheliums (HAEs) during single infections.**

A) Ratio of the trans-epithelial electrical resistance (TEER) over the starting TEER (T0 at day 0) during single infection of HAEs with SARS-CoV-2 (Omicron or D614G) or influenza A (H3N2 or H1N1). B) Percentage of viability over time compared to viability 24 h before infection, determined by a MTS assay. Results represent the mean ± SEM of 3-6 replicates from one or two independent experiments. C) Mean RNA copies per ml of 18S housekeeping gene in HAE lysates at 120 h p.i. ± SEM of 3-6 replicates from one or two independent experiments.

**S2 Fig. Interferon (IFN)-λ1 and λ2 production at 24 h after single infection with SARS-CoV-2 or IAV.**

Production of A) IFN-λ1 and B) IFN-λ2 proteins at the basolateral pole of nasal human airway epitheliums (HAEs) after single infections with SARS-CoV-2 (Omicron or D614G) and influenza A (H3N2 or H1N1), at 24 h post-infection. Non-infected HAEs are used as controls (NI). Results are expressed as the mean amount in pg per ml ± SEM of 3-4 replicates from one independent experiment.

**S3 Fig. Interferon-stimulated genes (ISGs) expression of uninfected nasal human airway epitheliums (HAEs) and during single infection with SARS-CoV-2 or IAV.**

A) Expression of four ISGs (OAS1, IFITM3, ISG15, MxA) in uninfected HAEs. B-D) Comparison of the expression of the different ISGs in HAEs infected with SARS-CoV-2 (Omicron or D614G) or IAV (H3N2 or H1N1) at 120 h p.i. Results are expressed as the mean of the ratio of ISG mRNAs over 18S housekeeping gene (both in copies per µL) ± SEM, calculated using 3-6 replicates from one or two independent experiments. *: p ≤ 0.05.

**S4 Fig. Viability of nasal human airway epitheliums (HAEs) during single infection with SARS-CoV-2 and IAV in the presence of ruxolitinib.**

A) Ratio of the trans-epithelial electrical resistance (TEER) over the starting TEER (T0 at day 0) and B) Percentage of viability (determined by a MTS assay) over time compared to viability 24 h before infection and during single infections of HAEs with SARS-CoV-2 (Omicron or D614G) or influenza A (H3N2 or H1N1), in the presence of ruxolitinib (ruxo). Results represent the mean ± SEM of 3-6 replicates from one or two independent experiments. C) Mean of the percentage of expression of 18S housekeeping gene in lysates of infected HAEs in the presence of ruxolitinib compared to the expression in untreated HAEs, at 120 h p.i. ± SEM of 3-6 replicates from one or two independent experiments. D) Inhibition of interferon-stimulated gene (ISG) mRNA expression by ruxolitinib in HAEs infected with SARS-CoV-2. Results are expressed as the mean inhibition percentage ± SEM of 9 replicates from two independent experiments.

**S5 Fig. Effect of BX795 on viral interference between Omicron and A/H3N2, and epithelium survival.**

A) Viral RNA loads in nasal human airway epitheliums (HAEs) infected with SARS-CoV-2 Omicron alone or in sequential coinfections (seq) 24 h after A/H3N2, in the presence of BX795. Results are expressed as the mean of the Log_10_ of viral RNA copies per ml ± SEM of 3 replicates from one experiment. **: p ≤ 0.01. B) Ratio of the trans-epithelial electrical resistance (TEER) over the starting TEER (T0 at day 0) and C) Percentage of viability (determined by a MTS assay) over time in HAEs infected with Omicron alone or sequentially with A/H3N2 and Omicron, compared to viability 24 h before infection, in the presence of BX795. Results represent the mean ± SEM of 3 replicates from one experiment.

## Notes

### Competing Interest Statement

The authors have declared no competing interest.

### Summary of Updates

We add a supplementary information file.

