## Supplementary material for "Viral interference between severe acute respiratory syndrome coronavirus 2 and influenza A viruses"

### Supporting information

**S1 Table. Sequences of primers and probes used for quantification of interferon-stimulated genes and a housekeeping gene by ddPCR.**

| | | Sequence | Reference |
|---|---|---|---|
| **OAS1** | Forward | 5'-GCA AAC AGG TCT GGG AGG-3' | This work |
| | Reverse | 5'-GTC AAT GGC ATG GTT GAT TTG C-3' | This work |
| | Probe | 5'-CAG TTC TGT TGC CAC TCT CTC TCC TG-3' | This work |
| **IFITM3** | Forward | 5'-ATC GTC ATC CCA GTG CTG AT-3' | (Cheemarla et al., 2021) |
| | Reverse | 5'-ATG GAA GTT GGA GTA CGT GG-3' | (Cheemarla et al., 2021) |
| | Probe | 5'-CAG GAG GCA TCA CTG AGG CCA G-3' | This work |
| **ISG15** | Forward | 5'-TGG ACA AAT GCG ACG AAC C-3' | This work |
| | Reverse | 5'-GGT CAG CCA GAA CAG GTC-3' | This work |
| | Probe | 5'-CTG GTG AGG AAT AAC AAG GGC CGC-3' | This work |
| **MxA** | Forward | 5'-GTC AGT TAC CAG GAC TAC GA-3' | This work |
| | Reverse | 5'- ATC TCC AGG GTG ATT AGC TC-3' | This work |
| | Probe | 5'-TTG AGA TTT CGG ATG CTT CAG AGG TAG-3' | This work |
| **18S** | Forward | 5'-GGA TGC GTG CAT TTA TCA G-3' | This work |
| | Reverse | 5'-AGT TGA TAG GGC AGA CGT TC-3' | This work |

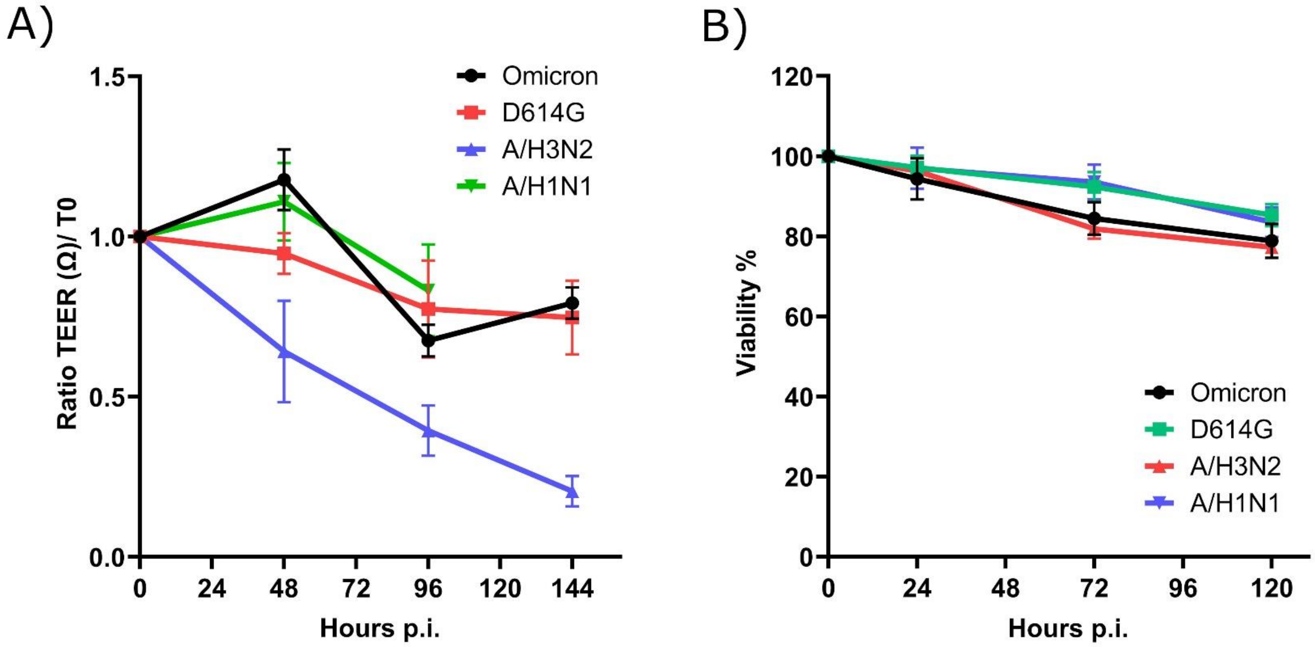


C)

| | 18S (Mean RNA copies/ml ± SEM) |
|---|---|
| SARS-CoV-2 Omicron | 202.7 ± 27.5 |
| SARS-CoV-2 D614G | 197.7 ± 9.8 |
| IAV H3N2 | 132.1 ± 17.9 |
| IAV H1N1 | 149.8 ± 8.5 |

**S1 Fig. Viability of nasal human airway epitheliums (HAEs) during single infections.**

A) Ratio of the trans-epithelial electrical resistance (TEER) over the starting TEER (T0 at day 0) during single infection of HAEs with SARS-CoV-2 (Omicron or D614G) or influenza A (H3N2 or H1N1). B) Percentage of viability over time compared to viability 24 h before infection, determined by a MTS assay. Results represent the mean ± SEM of 3-6 replicates from one or two independent experiments. C) Mean RNA copies per ml of 18S housekeeping gene in HAE lysates at 120 h p.i. ± SEM of 3-6 replicates from one or two independent experiments.

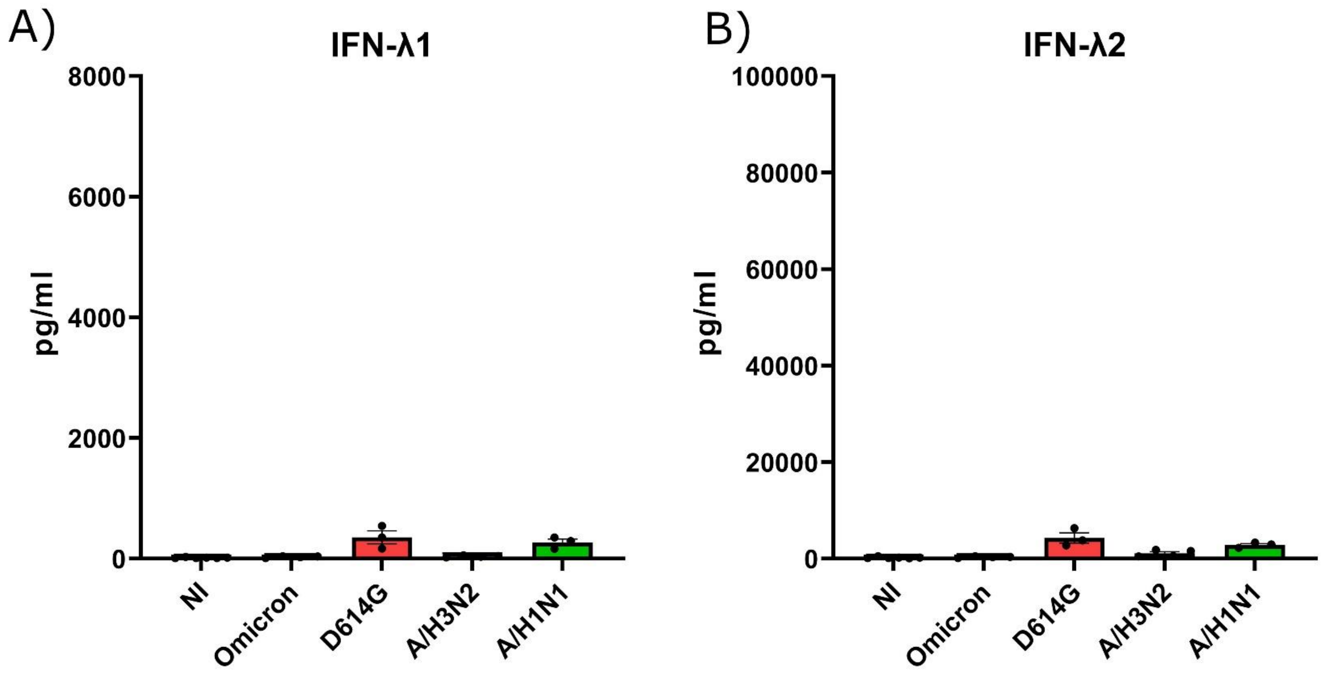


**S2 Fig. Interferon (IFN)-λ1 and λ2 production at 24 h after single infection with SARS-CoV-2 or IAV.**

Production of A) IFN-λ1 and B) IFN-λ2 proteins at the basolateral pole of nasal human airway epitheliums (HAEs) after single infections with SARS-CoV-2 (Omicron or D614G) and influenza A (H3N2 or H1N1), at 24 h post-infection. Non-infected HAEs are used as controls (NI). Results are expressed as the mean amount in pg per ml ± SEM of 3-4 replicates from one independent experiment.

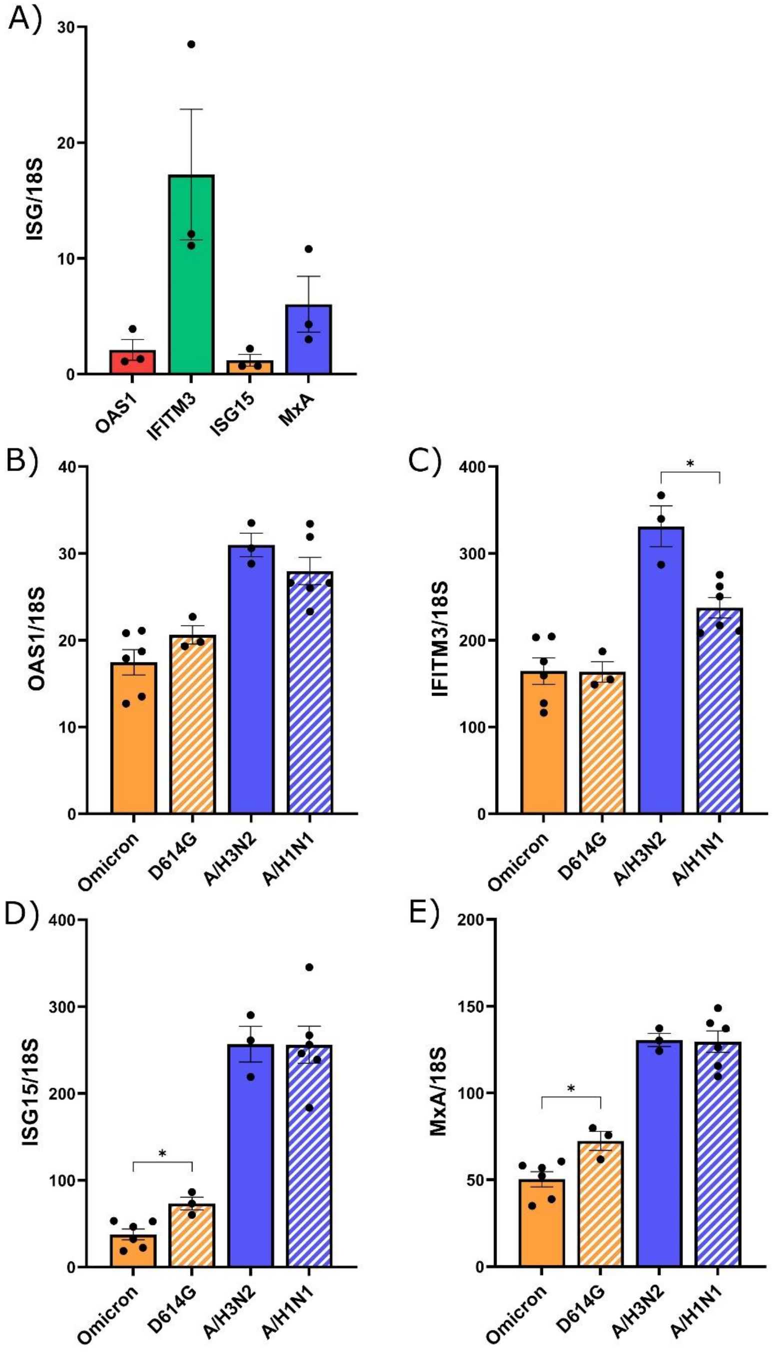


**S3 Fig. Interferon-stimulated genes (ISGs) expression of uninfected nasal human airway epitheliums (HAEs) and during single infection with SARS-CoV-2 or IAV.**

A) Expression of four ISGs (OAS1, IFITM3, ISG15, MxA) in uninfected HAEs. B-D) Comparison of the expression of the different ISGs in HAEs infected with SARS-CoV-2 (Omicron or D614G) or IAV (H3N2 or H1N1) at 120 h p.i. Results are expressed as the mean of the ratio of ISG mRNAs over 18S housekeeping gene (both in copies per µL) ± SEM, calculated using 3-6 replicates from one or two independent experiments. *: $p \leq 0.05$.

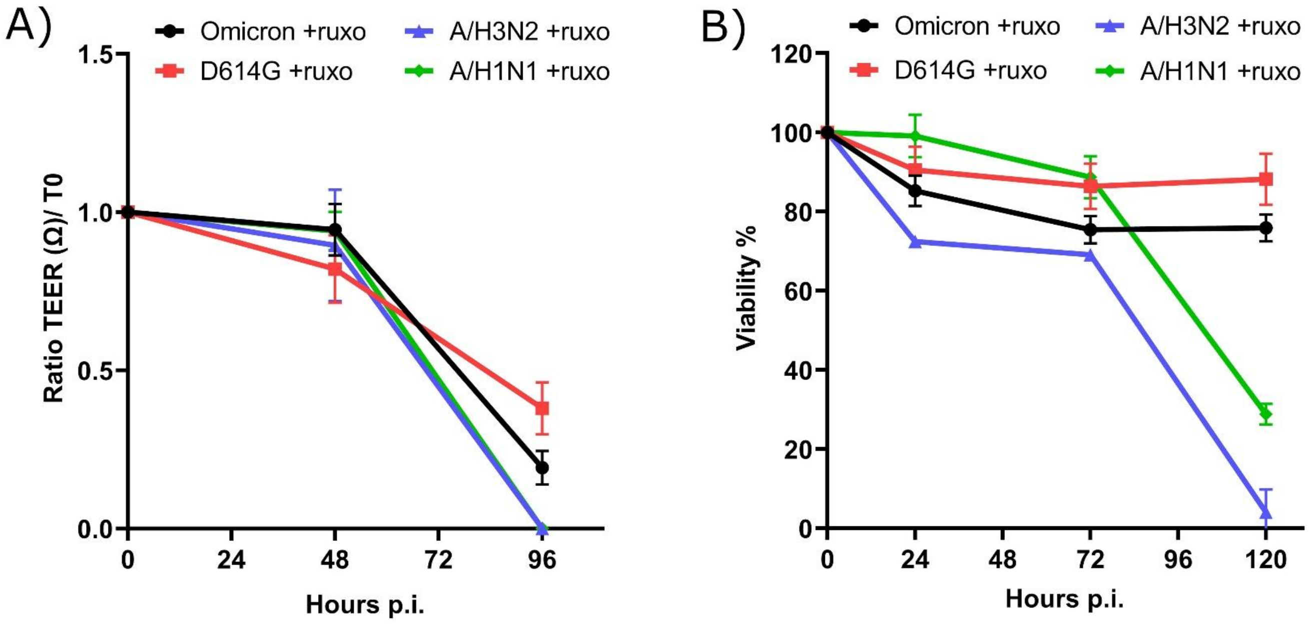


C)

| | **18S expression** +ruxolitinib /untreated (Mean percentage ± SEM) |
|---|---|
| SARS-CoV-2 Omicron | 77.1 ± 5.5 |
| SARS-CoV-2 D614G | 60.3 ± 1.2 |
| IAV H3N2 | 1.0 ± 0.7 |
| IAV H1N1 | 7.7 ± 0.4 |

D)

| ISGs | Mean inhibition % ± SEM |
|---|---|
| OAS1 | 79.2 ± 2.6 |
| IFITM3 | 90.3 ± 1.3 |
| ISG15 | 95.5 ± 0.6 |
| MxA | 92.8 ± 1.3 |

**S4 Fig. Viability of nasal human airway epitheliums (HAEs) during single infection with SARS-CoV-2 and IAV in the presence of ruxolitinib.**

A) Ratio of the trans-epithelial electrical resistance (TEER) over the starting TEER (T0 at day 0) and B) Percentage of viability (determined by a MTS assay) over time compared to viability 24 h before infection and during single infections of HAEs with SARS-CoV-2 (Omicron or D614G) or influenza A (H3N2 or H1N1), in the presence of ruxolitinib (ruxo). Results represent the mean ± SEM of 3-6 replicates from one or two independent experiments. C) Mean of the percentage of expression of 18S housekeeping gene in lysates of infected HAEs in the presence of ruxolitinib compared to the expression in untreated HAEs, at 120 h p.i. ± SEM of 3-6 replicates from one or two independent experiments. D) Inhibition of interferon-stimulated gene (ISG) mRNA expression by ruxolitinib in HAEs infected with SARS-CoV-2. Results are expressed as the mean inhibition percentage ± SEM of 9 replicates from two independent experiments.

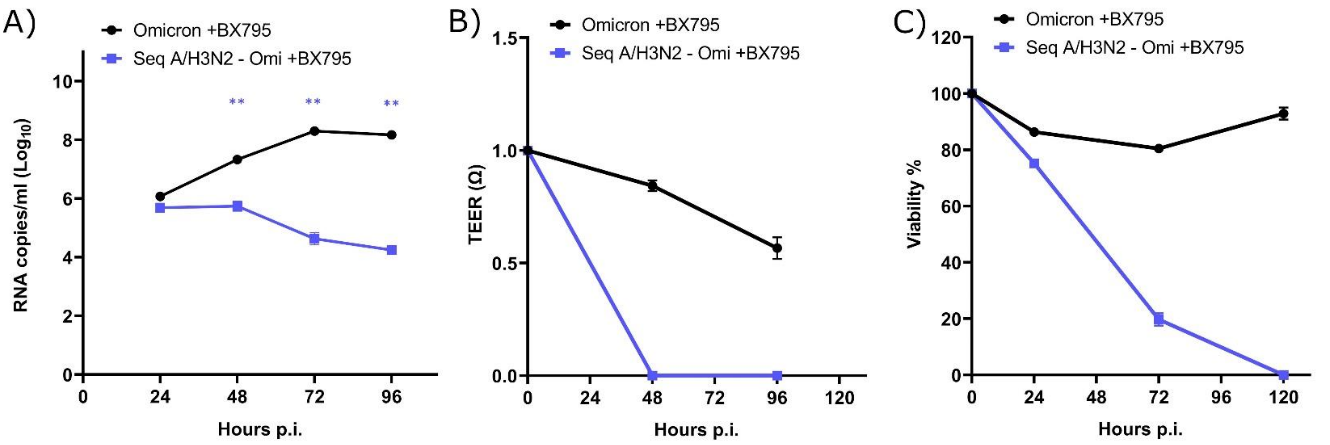


**S5 Fig. Effect of BX795 on viral interference between Omicron and A/H3N2, and epithelium survival.**

A) Viral RNA loads in nasal human airway epitheliums (HAEs) infected with SARS-CoV-2 Omicron alone or in sequential coinfections (seq) 24 h after A/H3N2, in the presence of BX795. Results are expressed as the mean of the $Log_{10}$ of viral RNA copies per ml ± SEM of 3 replicates from one experiment. **: $p \leq 0.01$. B) Ratio of the trans-epithelial electrical resistance (TEER) over the starting TEER (T0 at day 0) and C) Percentage of viability (determined by a MTS assay) over time in HAEs infected with Omicron alone or sequentially with A/H3N2 and Omicron, compared to viability 24 h before infection, in the presence of BX795. Results represent the mean ± SEM of 3 replicates from one experiment.
